# Nanoligomers targeting NF-κB and NLRP3 reduce neuroinflammation and improve cognitive function with aging and tauopathy

**DOI:** 10.1101/2024.02.03.578493

**Authors:** Devin Wahl, Sydney J. Risen, Shelby C. Osburn, Tobias Emge, Sadhana Sharma, Vincenzo S. Gilberto, Anushree Chatterjee, Prashant Nagpal, Julie A. Moreno, Thomas J. LaRocca

## Abstract

Neuroinflammation contributes to impaired cognitive function in brain aging and neurodegenerative disorders like Alzheimer’s disease, which is characterized by the aggregation of pathological tau. One major driver of both age- and tau-associated neuroinflammation is the NF-κB and NLRP3 signaling axis. However, current treatments targeting NF-κB or NLRP3 may have adverse/systemic effects, and most have not been clinically translatable. In this study, we tested the efficacy of a novel, nucleic acid therapeutic (Nanoligomer) cocktail specifically targeting both NF-κB and NLRP3 in the brain for reducing neuroinflammation and improving cognitive function in old (aged 19 months) wildtype mice, and in rTg4510 tau pathology mice (aged 2 months). We found that 4 weeks of NF-κB/NLRP3-targeting Nanoligomer treatment strongly reduced neuro-inflammatory cytokine profiles in the brain and improved cognitive-behavioral function in both old and rTg4510 mice. These effects of NF-κB/NLRP3-targeting Nanoligomers were also associated with reduced glial cell activation and pathology, favorable changes in transcriptome signatures of glia-associated inflammation (reduced) and neuronal health (increased), and positive systemic effects. Collectively, our results provide a basis for future translational studies targeting both NF-κB and NLRP3 in the brain, perhaps using Nanoligomers, to inhibit neuroinflammation and improve cognitive function with aging and neurodegeneration.

## 1. INTRODUCTION

Aging is associated with declines in cognitive function, and it is the primary risk factor for most neurodegenerative diseases, including Alzheimer’s disease (AD). These declines in cognitive function are closely linked with the accumulation of pathological tau, a hallmark of AD that also increases with aging [1, 2]. One key “macro-mechanism” that contributes to both brain aging and tau pathology is neuroinflammation, which is characterized by innate immune signaling, glial cell activation, reduced neuronal health and the release of neurotoxic, pro-inflammatory cytokines [3]. In fact, evidence suggests that neuroinflammation may occur early in brain aging and both precede and be exacerbated by tauopathy [4]. As such, identifying strategies to reduce neuroinflammation in the context of aging and tauopathy is an important biomedical research goal [5].

The transcription factor Nuclear Factor Kappa B (NF-κB) and inflammasome member NLR Family Pyrin Domain Containing 3 (NLRP3) are key drivers of age- and tau-associated neuroinflammation, and both play central roles in adverse processes that drive brain aging and AD (e.g., immune activation and pro-inflammatory signaling) [6, 7]. The roles of these proteins in brain aging and neurodegenerative disease are multifaceted, and NF-κB in particular may also be neuroprotective (e.g., by regulating synaptic plasticity and neurotrophic factors) [8]. However, a preponderance of recent studies have demonstrated that the inhibition of either NF-κB or NLRP3 alone reduces immune activation/pro-inflammatory cytokine release in the brain and improves cognitive function [9–11], suggesting that the NF-κB/NLRP3 signaling axis may be a promising therapeutic target. Despite these findings, current treatments that target NF-κB and NLRP3 may have off-target effects (e.g., by inhibiting activity of other enzymes) [12] and/or poor biodistribution, and many systemic anti-inflammatory treatments (e.g., immunosuppressive steroids) do not cross the blood-brain barrier. To address these concerns, our team recently developed a novel Nanoligomer (nucleic acid therapeutic) cocktail that targets both NF-κB and NLRP3 DNA, to inhibit transcription, and mRNA, to inhibit translation (**Figure 1A**).

**Figure 1.**
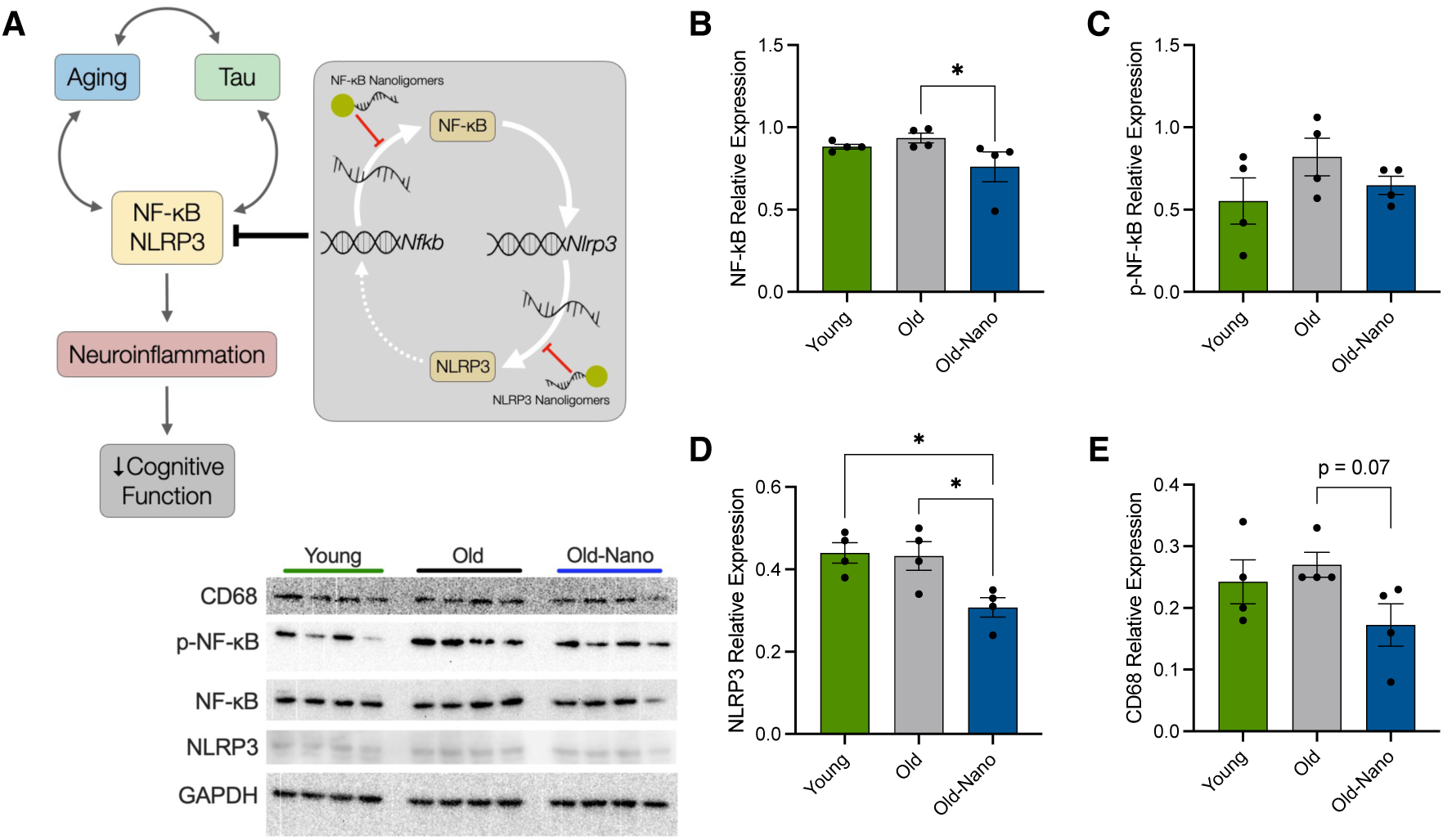
Short-term Nanoligomer treatment downregulates NF-κB and NLRP3 in the brain. **(A)** NF-κB and NLRP3 interact to drive inflammation, but Nanoligomers that block their translation may reduce age- and tau-associated neuroinflammation. **(B-E)** Western blot data showing protein levels of NF-κB, phosphorylated (p)-NF-κB, NLRP3 and Cluster of Differentiation 68 (CD68) in the hippocampus of young (8 months) and older (19 months) wildtype mice, and older mice 16 h after treatment with a NF-κB/NLRP3-targeting Nanoligomer cocktail. Raw immunoblots shown at bottom left. Data analyzed via non-parametric one-way ANOVA and group differences determined via Kruskal-Wallace post-hoc test. N = 4/group; *p < 0.05.

Nanoligomers comprise an antisense peptide nucleic acid (PNA) conjugated to a gold nanoparticle for improved delivery [13–18], and this particular Nanoligomer cocktail was developed via in-depth screening studies for different targets to reduce neuroinflammation, in which NF-κB emerged as the top single target and NLRP3 showed a synergistic interaction [16, 19]. Our recent reports demonstrate that this NF-κB/NLRP3 Nanoligomer cocktail is highly specific, crosses the blood-brain barrier, is bioavailable across various brain regions [18, 20], downregulates its targets both at the RNA and protein level, and inhibits neuroinflammation *in vitro* and *in vivo* [15, 16, 18–20], yet its influence on cognitive function in the context of brain aging and tauopathy has not been thoroughly explored.

In the present study, we evaluated the efficacy of targeting both NF-κB and NLRP3 in brain aging and tauopathy by treating both old wildtype mice and transgenic rTg4510 mice (a model of tauopathy commonly used in neurodegeneration research) with our Nanoligomer cocktail. We found that this treatment: 1) improved cognitive function and broadly reduced neuroinflammation in both models; 2) reduced tau pathology and glial cell activation while conferring neuroprotection; 3) reversed transcriptome changes common to brain aging and tauopathy; and 4) exerted favorable systemic effects. Our results show that selective targeting of both NF-κB and NLRP3 is a potent strategy for reducing neuroinflammation, and they provide a basis for similar studies in larger animal models and/or pilot clinical trials.

## 2. RESULTS

### 2.1. Short-term Nanoligomer treatment downregulates NF-κB and NLRP3 in the hippocampus

NF-κB and NLRP3 are central mediators of age-related inflammation (i.e., “inflammaging”), neuroinflammation, innate immune responses and cytokine release, including in the brain [6, 7, 21, 22]. Our published data show that NF-κB/NLRP3-targeting Nanoligomers downregulate these targets in multiple brain regions [18, 20].

However, to confirm this in the present study, we first treated older (aged 19 months) male and female C57Bl/6J mice with the same Nanoligomer cocktail or saline control (1x intraperitoneal injection; 150 mg/kg body weight). Younger (aged 8 months) male and female mice, as a reference group, also received a saline injection. After 16 hours, mice were euthanized and immunoblots were performed on isolated whole hippocampus. We observed significant reductions in both NF-κB and NLRP3 in old treated vs. untreated mice (**Figure 1B,D**), and a modest but non-significant increase in phosphorylated (activated) NF-κB in older vs. young control animals that was absent in older treated animals (**Figure 1C**). These effects of acute Nanoligomer administration on central mediators of neuroinflammation and innate immune responses are consistent with our reports showing that the drug crosses the blood-brain barrier and engages its targets [13, 14, 16, 18, 20], suggesting the potential for protective effects on mediators of neuroinflammation downstream of NF-κB/NLRP3 (e.g., reactive glia, pro-inflammatory gene induction and cytokine release) [23, 24].

Consistent with this idea, even with the short duration of this treatment, we also found a nearly significant trend for reduced expression of Cluster of Differentiation 68 (CD68), a marker of pro-inflammatory microglia in Nanoligomer-treated old mice (**Figure 1D**).

### 2.2. Long-term Nanoligomer treatment improves cognitive function in old wildtype and tauopathy mice

Although NF-κB and NLRP3 play complex roles in brain health [8], most data suggest that inhibiting these proteins individually improves cognitive function [25, 26]. Therefore, given our finding that acute Nanoligomer treatment reduced NF-κB and NLRP3 in the hippocampus (a key area of the brain involved in memory and cognitive function) and the central role of neuroinflammation in cognitive dysfunction with brain aging and AD [27], we next tested the hypothesis that long-term Nanoligomer treatment (150 mg/kg body weight; 3x week for one month) would improve cognitive function in old C57Bl/6J male and female mice. In addition, because tauopathy (a key hallmark of AD) is associated with NF-κB/NLRP3 activation [28] and cognitive dysfunction [29], we also treated 2 month-old male and female rTg4510 mice, a common model of tauopathy based on expression of human P301L tau in the forebrain, using the same approach. Importantly, we did not note any adverse effects with this treatment, and all body weights remained stable. After the treatment, we measured short-term recognition memory via Novel Object Recognition testing and anxiety-like behavior via Elevated Plus Maze testing. In Novel Object Recognition testing, we found that treated older and tauopathy mice performed significantly better (as measured by recognition indices) than untreated controls, and similarly to younger mice and littermate controls, respectively (**Figure 2A**). In the Elevated Plus Maze test, we observed a non-significant trend for treated older and tauopathy mice to spend more time in open maze arms than untreated mice, reflecting reduced anxiety (**Figure 2B**). Finally, in tests of grip strength, which we measured because of its relationship to cognitive function and overall health [30], we found a slight but non-significant trend for improvements in old treated mice, although these effects were not noted in treated tauopathy mice, and among all animals we observed no differences in frailty, which is closely associated with cognitive decline [31] (**Supplementary Figure S1**). Future, more comprehensive studies are needed to assess how NF-κB/NLRP3-targeting Nanoligomer treatment affects other cognitive and peripheral functional outcomes (e.g., spatial memory, treadmill endurance, wire hang, rotarod performance). However, these results show that long-term Nanoligomer treatment improves several domains of cognitive-behavioral function in old, non-transgenic mice, and in tauopathy model mice, without overt adverse effects on neuromuscular function/health.

**Figure 2.**
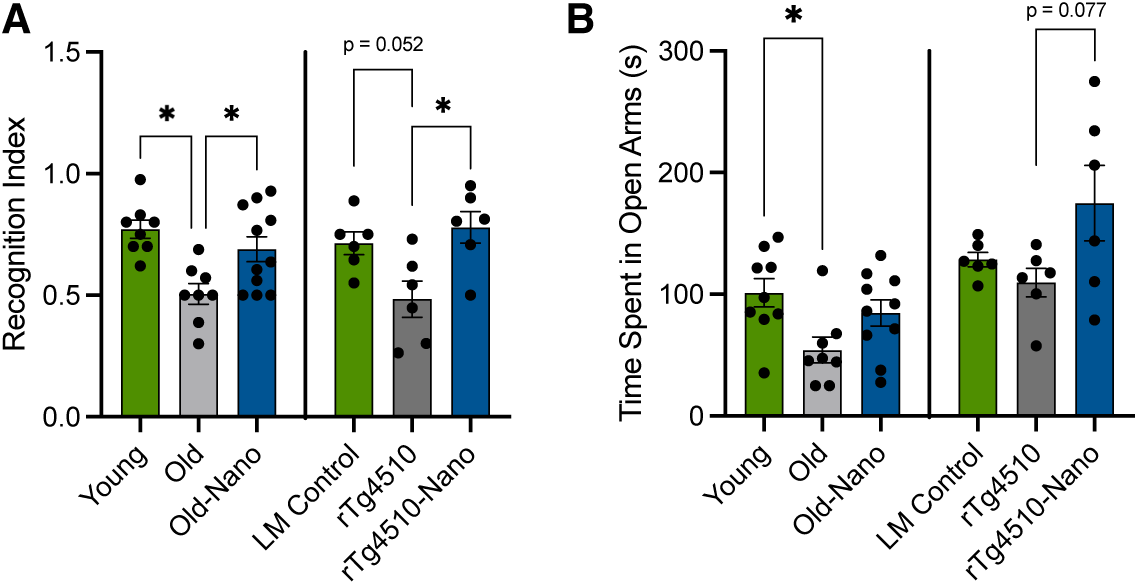
Long-term treatment with Nanoligomers targeting NF-κB and NLRP3 improves cognitive-behavioral function in old wildtype and tauopathy mice. **(A)** Novel object recognition index (short-term recognition memory) quantified as time exploring the novel object versus total object exploration time in young, old and old Nanoligomer-treated wildtype mice, as well as littermate (LM) controls, rTg4510 tauopathy and rTg4510 Nanoligomer treated mice. **(B)** Time spent in open arms during Elevated Plus Maze testing, a reflection of anxiety-like behavior, in the same mice. Data analyzed via one-way ANOVA and group differences determined via Tukey’s post-hoc multiple comparison test. N = 6-11/group; *p < 0.05.

### 2.3. Broad Nanoligomer-induced reductions in neuroinflammation correlate with cognitive function

The activation of NF-κB and NLRP3 via aging or tauopathy induces the production of many pro-inflammatory cytokines [32]. Therefore, we performed multiplex ELISAs on brains from the same mice (long-term Nanoligomer-treated) described above, to assess levels of key cytokines involved in neuroinflammation and innate immune responses. Principal component analyses (PCA) revealed that aging and tauopathy were associated with robust and broad increases in many cytokines, whereas cytokine levels in treated old and tauopathy animals were similar to those in young mice and littermate controls, respectively (**Figure 3A and 3B**, left panels). At the individual cytokine level, several important pro-inflammatory and immune-activating cytokines that increased with aging and tauopathy included Interleukin 5 (IL-5) and Macrophage-Inflammatory Protein 2 (MIP-2α), but these effects were reduced with Nanoligomer treatment. Similar trends were noted for most Interleukins (ILs), Monocyte Chemoattractant Proteins (MCPs), and Interferons (IFNs) (**Supplementary Figure S2**). Additionally, we found that cytokine levels (global/PC-based and individual) were significantly inversely correlated with short-term recognition memory (**Figure 3A,B** center and right panels), which is consistent with existing evidence that increased cytokine levels in the brain are associated with cognitive decline [33]. Together, these data show that long-term NF-κB/NLRP3-targeting Nanoligomer treatment broadly reduces neuroinflammation, including many cytokines that are directly modulated by NF-κB and NLRP3 [22], and that these changes may contribute to improvements in cognitive function in old wildtype and tauopathy model mice.

**Figure 3.**
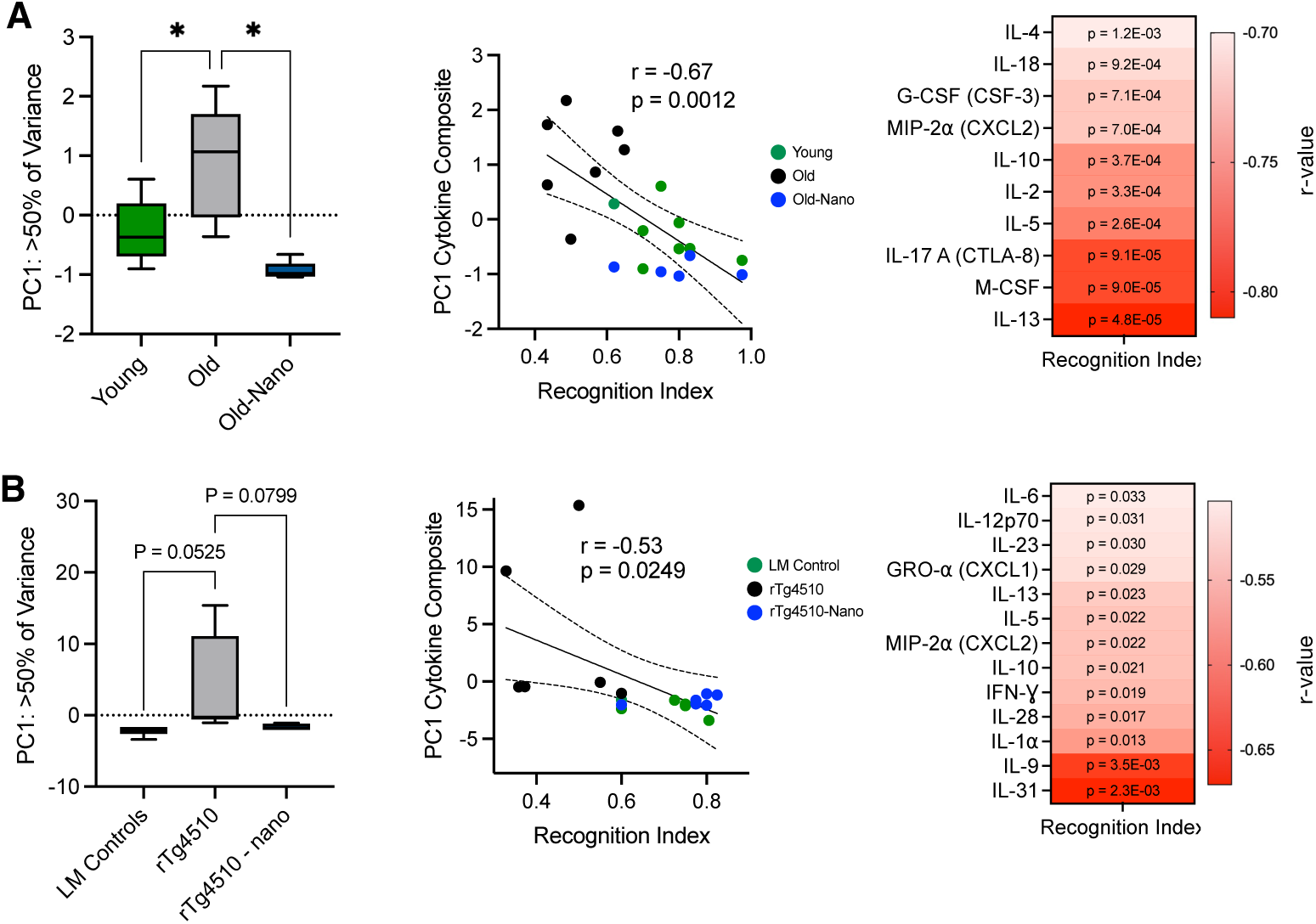
Broad reductions in neuroinflammation with Nanoligomer treatment are related to greater cognitive-behavioral function in old wildtype and tauopathy mice. **(A)** Principal Component 1 (PC1) from PC analysis of 36 cytokines in hippocampus of young, old and old Nanoligomer-treated wildtype mice (left panel). Correlation of PC1 values with Novel Object Recognition Index scores in the same mice (middle panel) and heatmap showing magnitude of individual cytokine correlations with Recognition Index (right panel). **(B)** The same PC and cytokine analyses in littermate (LM) controls, rTg4510 tauopathy and rTg4510 Nanoligomer-treated mice. Correlations determined as simple Pearson correlation coefficients (r), and all analyses N = 6-11/group; *p < 0.05.

### 2.4. Reductions in phosphorylated tau with Nanoligomer treatment

Importantly, tau pathology can both activate and be exacerbated by NF-κB and NLRP3 [34–36], driving cycles of inflammation-mediated tau propagation and toxicity [37], and phosphorylated (activated) tau is closely associated with cognitive dysfunction [38]. To characterize the effects of Nanoligomer treatment on tau, we performed immunohistochemistry for two insoluble tau residues (phosphorylated at threonine-181 and threonine-231) that have been associated with AD pathology and progression [39, 40]. We found that these tau species increased in hippocampus and cortex of rTg4510 mice, similar to what others have reported in other tauopathy models [41, 42], and that this effect was attenuated with Nanoligomer treatment (**Figure 4A**). We also performed immunoblots for tau, including for phosphorylated species more typical of “later” pathology, and we found that long-term Nanoligomer treatment was associated with modest reductions (p = 0.05) in both total and phosphorylated (at serine 396) tau using this approach (**Figure 4B**).

**Figure 4.**
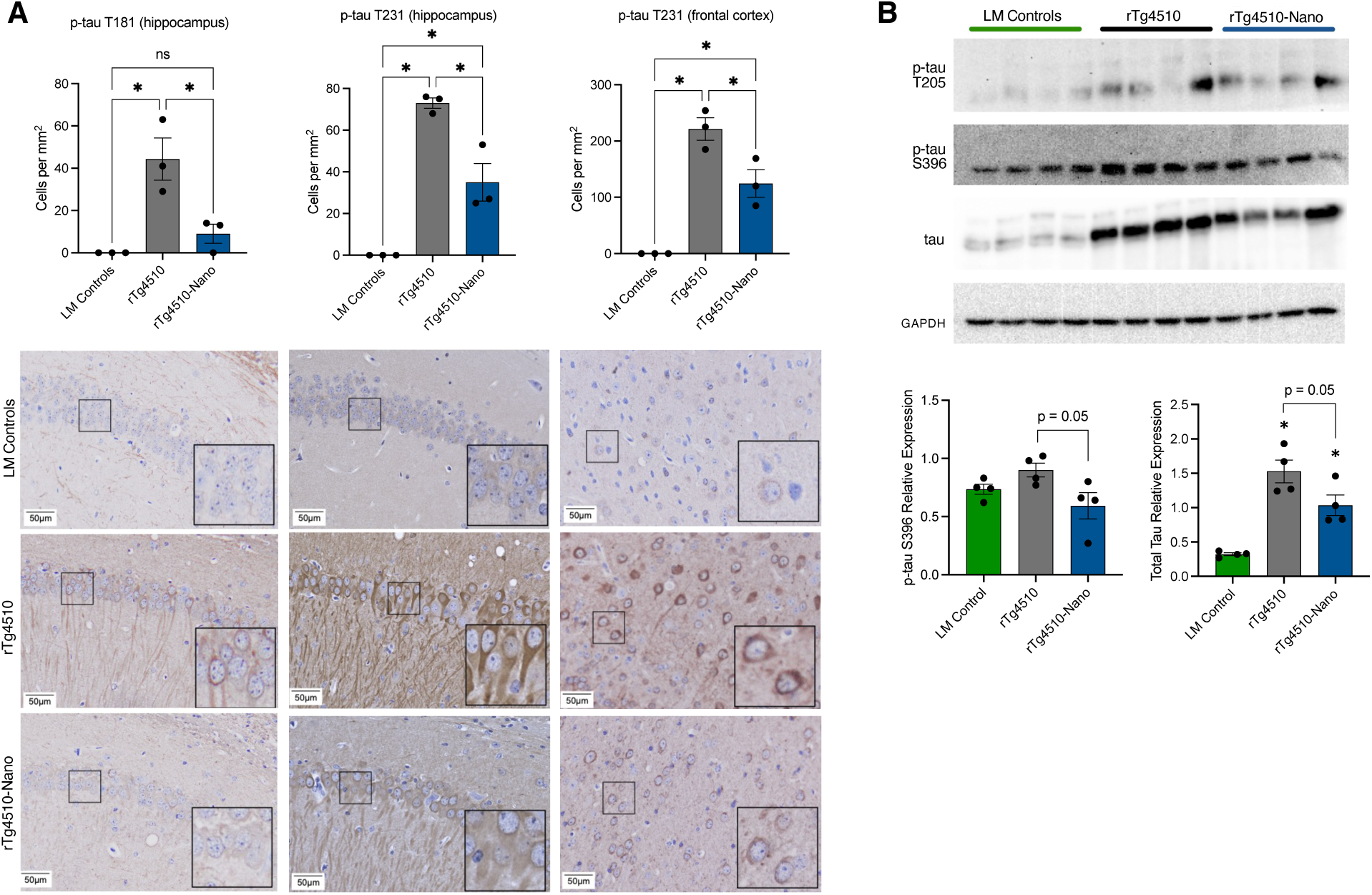
Reductions in phosphorylated tau in the hippocampus with Nanoligomer treatment. **(A)** Quantification and representative immunohistochemistry images of phosphorylated tau (p-threonine-181 and p-threonine 231) in hippocampal and frontal cortex sections from littermate (LM) controls, rTg4510 tauopathy and rTg4510 Nanoligomer-treated mice. Insets in lower right corner of images are 2x magnification of representative cell bodies indicated in smaller boxes. Note reduction in p-tau staining density with Nanoligomer treatment. **(B)** Raw immunoblots and quantification of total and phosphorylated tau species (p-threonine 205 and p-serine 396) in the same mice. Data analyzed via one-way ANOVA and group differences determined via Tukey’s post-hoc multiple comparison test. Immunohistochemistry, N = 3/group; immunoblots, N = 4/group; *p < 0.05.

Finally, in these same mice, additional immunohistochemistry analyses showed that Nanoligomer treatment and the reduced tau burden we observed were associated with: 1) neuroprotection, as reflected by greater numbers of Fox-3 (NeuN)-positive cells in the CA1 and CA2 regions of the hippocampus (**Figure 5A**); and 2) fewer potentially reactive glia, as reflected by reductions in Glial Fibrillary Acidic Protein (GFAP)-positive astrocytes and a non-significant trend for reductions in Ionized Calcium-Binding Adapter molecule 1 (IBA1)- positive microglia (**Figure 5B**). Although GFAP and IBA1 counts do not reflect neuroinflammation per se, these results suggest that NF-κB/NLRP3-targeting Nanoligomer treatment may suppress tau pathology and confer neuroprotection, and that this effect may involve changes in glial cell-associated processes.

**Figure 5.**
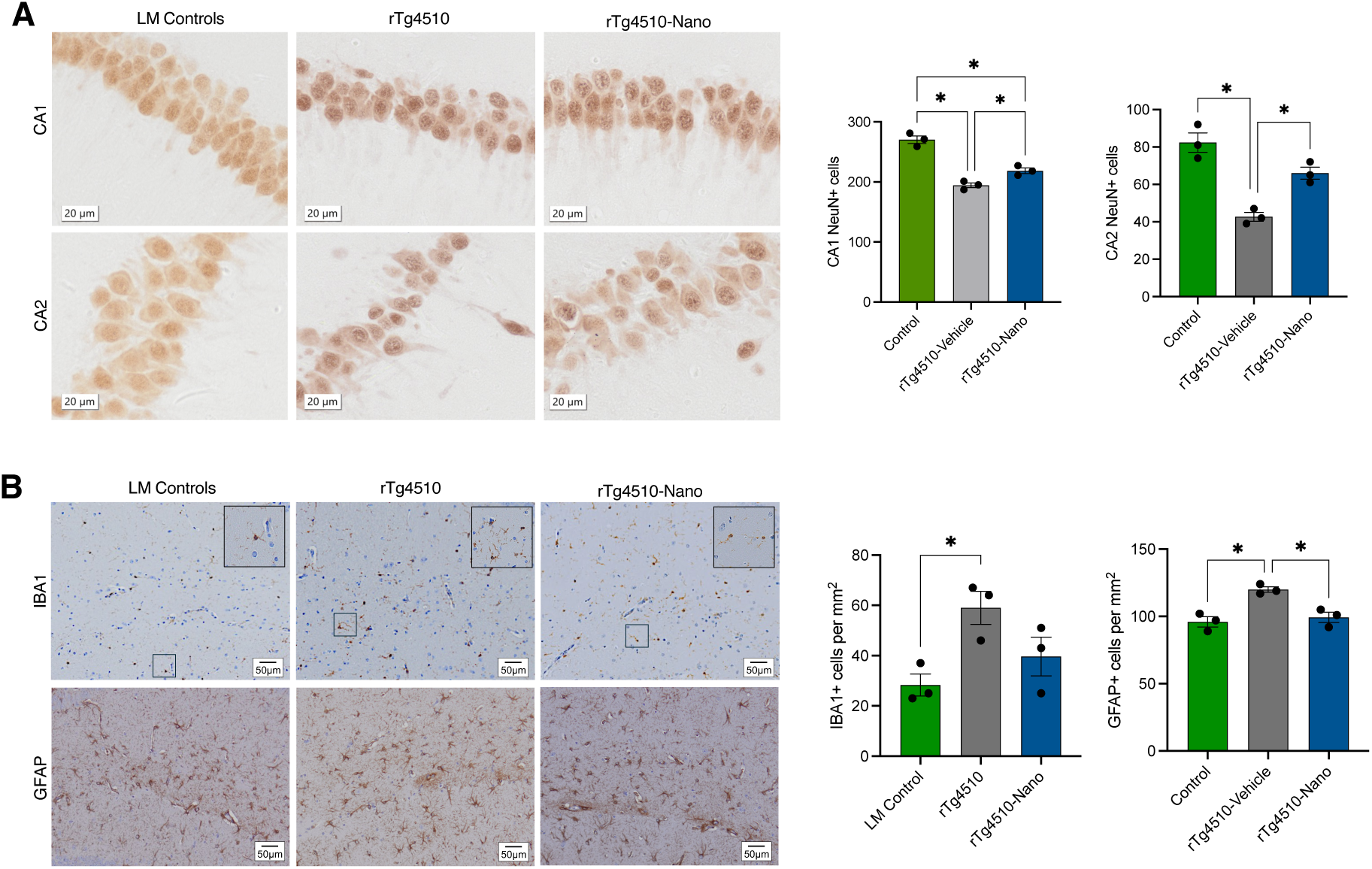
Changes in glial cell and neuronal numbers with Nanoligomer treatment in tauopathy mice. **(A)** Representative immunohistochemistry images (left) and counts (right) for cells positive for Fox-3 (NeuN) neuronal marker in CA1 and CA2 regions of the hippocampus in littermate (LM) controls, rTg4510 tauopathy and rTg4510 Nanoligomer-treated mice. **(B)** Immunohistochemistry images (left) and counts (right) for cells positive for Ionized Calcium-Binding Adapter molecule 1 (IBA1) and Glial Fibrillary Acidic Protein (GFAP) in the same mice. Data analyzed via one-way ANOVA and group differences determined via Tukey’s post-hoc multiple comparison test. N = 3/group; *p < 0.05.

### 2.5. Reversal of age-related glial cell activation with Nanoligomer treatment

Although aged wildtype mice do not develop tau pathology, they do develop gliosis like humans. Indeed, in human aging/AD, glial cells (astrocytes and microglia) become immune-activated, secreting cytokines that impair neuronal and cognitive function [43]. This neuroinflammation is also associated with morphological changes [44, 45], including alterations in process length [46] and branching [47, 48], which have detrimental effects on neurons (e.g., synaptic dysfunction). To evaluate the effects of NF-κB/NLRP3-targeting Nanoligomer treatment on astrocyte and microglia activation/morphology with aging, we performed immunofluorescence staining on cortex tissue from old, old treated, and young mice. Consistent with previous findings showing that astrocyte activation increases with aging [44, 49], we found evidence of age-related increases in GFAP [50], but this effect was less pronounced in old treated mice (**Figure 6A**). Similar to previous studies [51, 52], we also found that aging was associated with reduced astrocyte process length and branch number (**Figure 6B**). However, these effects were reversed with Nanoligomer treatment. We also found age-related increases in the activation of microglia, as reflected by increases in IBA1 (**Figure 6C**). This effect of aging on microglia was not altered by Nanoligomer treatment, although we did observe changes in microglial process length and branch number in old animals that were attenuated with treatment (**Figure 6D**). Finally, similar to our observations in rTg4510 mice, using immunohistochemistry we found that all of these glial cell changes were associated with a reduction in NeuN+ cells in old mice that was absent in treated animals, although only in the CA2 region of the hippocampus (**Figure 6E**). Collectively, these results suggest that the beneficial effects of Nanoligomers targeting NF-κB/NLRP3 in old mice may be due in part to reducing age-related changes in glial cells and the preservation of neurons in the hippocampus.

**Figure 6.**
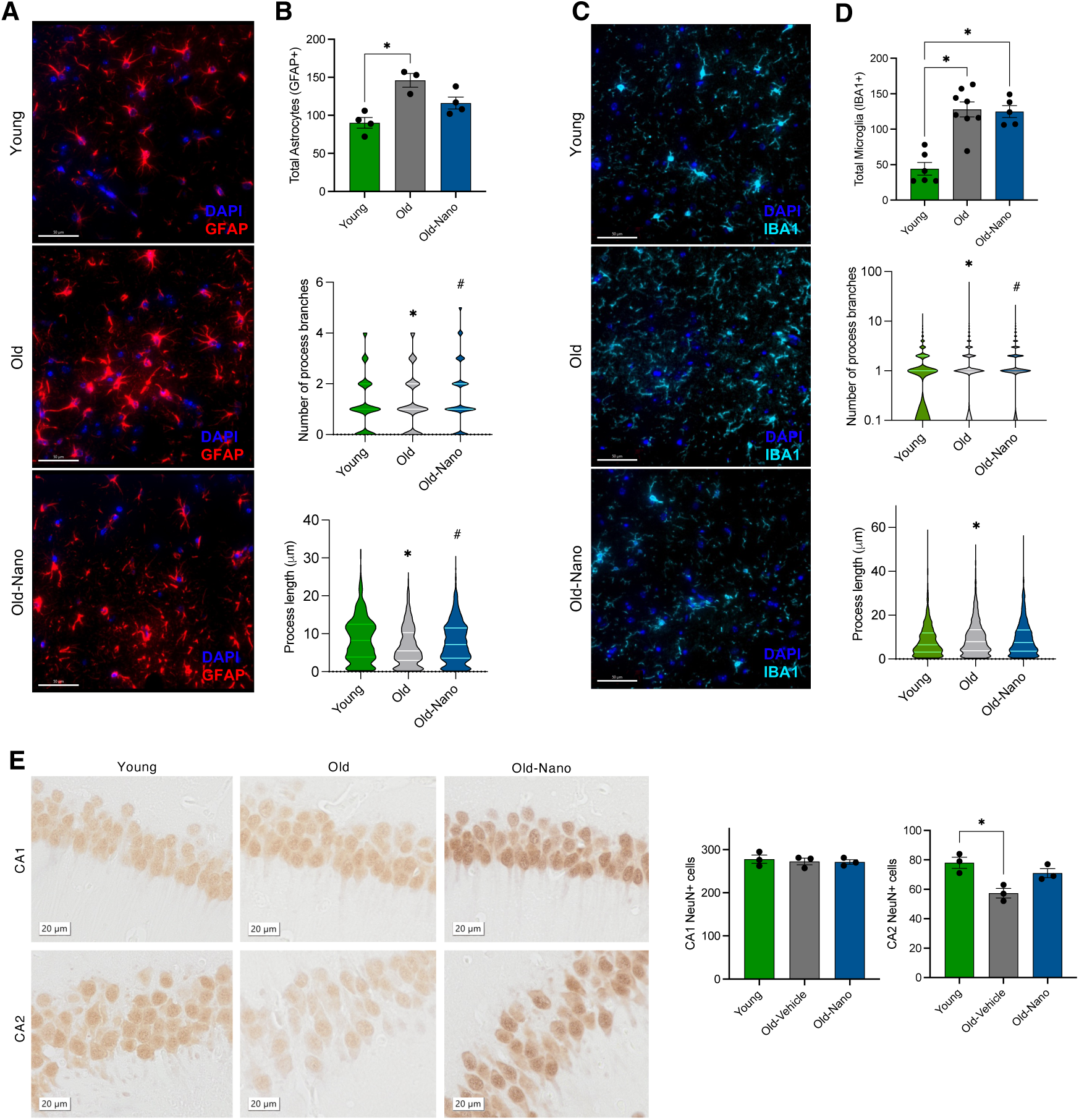
Reversal of age-related cellular changes in astrocytes and microglia with Nanoligomer treatment. **(A)** Immunofluorescence images of Glial Fibrillary Acidic Protein (GFAP) staining in hippocampus sections from young, old and old Nanoligomer-treated wildtype mice. Note general increase in GFAP/altered astrocyte morphology in old mice that is reversed with Nanoligomer treatment. **(B)** Quantification of GFAP+ cells and process length and branch number. **(C)** Immunofluorescence images of Ionized Calcium-Binding Adapter molecule 1 (IBA1) staining in the same mice. Note general increase in IBA1/altered microglial morphology in old mice that is reversed with treatment. **(D)** Quantification of IBA1+ cells and process length and branch number. **(E)** Immunohistochemistry counts for Fox-3 (NeuN)-positive cells (neurons) in CA1 and CA2 regions of the hippocampus in the same mice. *p < 0.05 vs. young; #p < 0.01 vs. old control.

### 2.6. Transcriptome signatures of inflammation and reduced neuronal health are reversed with Nanoligomer treatment

Perturbations in the transcriptome affect downstream biological processes that mediate brain aging/AD [53]. To provide broad insight into how Nanoligomer treatment affected genes/pathways most related to aging and tauopathy, we performed RNA-seq on the hippocampus. We found that aging and tauopathy were associated with gene expression changes (**Figure 7A**, top rows of heatmaps) that were largely reversed in terms of relative expression with Nanoligomer treatment (**Figure 7A**, bottom rows of heatmaps). For example, one of the top transcripts that increased with aging but decreased with Nanoligomer treatment was S100 calcium-binding protein A8 (*S100A8*), which activates NF-κB and induces microglial activation [54]. One of the top transcripts that increased with tauopathy but decreased with Nanoligomer treatment was Interleukin 17 Receptor E (*IL17RE*), which is also associated with the activation of NF-κB [55] and cognitive dysfunction [56].

**Figure 7.**
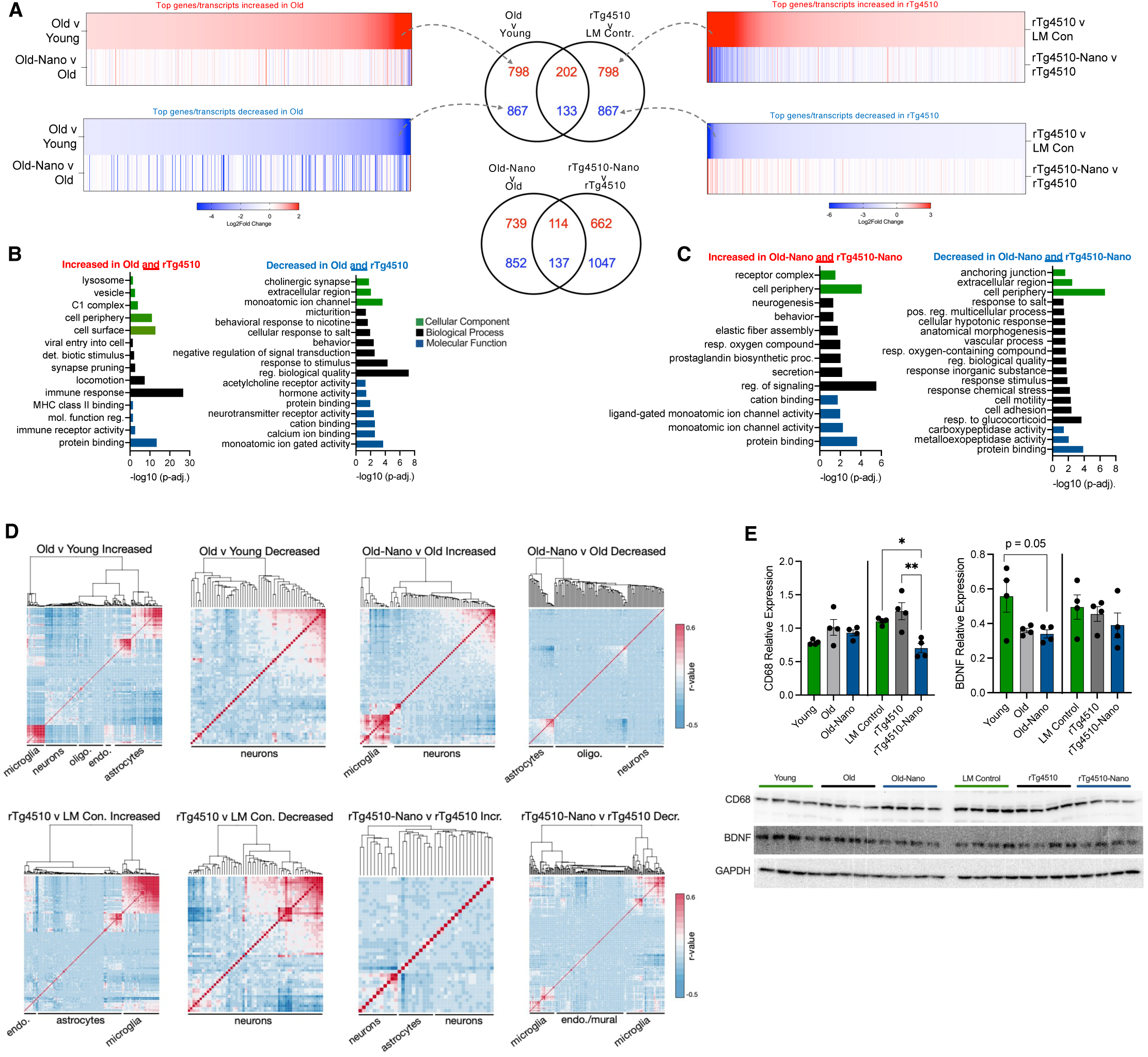
Transcriptome signatures of inflammation and reduced neuronal health with aging and tauopathy are reversed with Nanoligomer treatment. **(A)** Heatmaps showing Log2Fold differences for the top 1000 increased and decreased transcripts in old vs. young control mice and rTg4510 tauopathy mice vs. littermate (LM) controls (top heatmap rows), and Log2Fold differences for the same genes/transcripts in treated old and rTg4510 mice vs. their untreated counterparts (bottom heatmap rows). Venn diagrams in center show genes/transcripts that were increased or decreased in both old and rTg4510 mice, and genes/transcripts that were reversed in terms of relative expression in both treated old and treated rTg4510 mice. **(B)** Gene ontology analyses of genes/transcripts that were increased or decreased in both old and rTg4510 mice. **(C)** Gene ontology analyses of genes/transcripts that were increased or decreased in both treated old and treated rTg4510 mice. **(D)** Heatmaps showing hierarchical clustering and correlations between gene/transcript expression levels in bulk RNA-seq data from the present study and a single-cell RNA-seq dataset. Cell types associated with major gene expression clusters annotated; note glial cells associated with increased clusters and neurons associated with decreased clusters in both old and rTg4510 mice, and partial reversal of these patterns in heatmaps showing Nanoligomer treatment. **(E)** Quantifications and raw immunoblots for Cluster of Differentiation 68 (CD68) and Brain-Derived Neurotrophic Factor (BDNF) in animals from all groups. Gene expression differences analyzed using DESeq2 algorithms and immunoblot data analyzed via one-way ANOVA with group differences determined via Tukey’s post-hoc multiple comparison test. N = 4/group for all analyses; *p < 0.05.

Given the above observations and because aging and tau pathophysiology intersect [57], to understand how Nanoligomer treatment affected genes/pathways common to both aging and tauopathy, we compared the top 1000 most increased/decreased transcripts common to both aging and tauopathy, and those same transcripts in old and tauopathy-treated mice (**Figure 7A**, top Venn diagrams). Gene ontology (GO) analyses revealed that increased genes common to both aging and tauopathy reflected pathways associated with inflammation, immune responses and cognitive dysfunction, including immune receptor activity, major histocompatibility complex (MHC) binding, and synapse pruning (**Figure 7B**, left). Decreased genes/transcripts common to both aging and tauopathy comprised pathways associated with neurotransmitter receptor activity and signal transduction (**Figure 7B**, right). However, these patterns were generally reversed with treatment. That is, among the transcripts that changed in both treated old and treated tauopathy mice (**Figure 7A**, bottom Venn diagrams), those that were increased reflected pathways involved in brain aging and neuronal/cognitive function including ion channel activity, neuronal signaling, and neurogenesis (**Figure 7C**, left), whereas common decreased transcripts with treatment in aged and tauopathy mice reflected pathways associated with inflammation and immune responses, including cell adhesion and stress response (**Figure 7C**, right).

To determine which cell types might have contributed most to the observed transcriptome patterns, we performed a deconvolution analysis (which intersects bulk RNA-seq data with single-cell RNA-seq data to identify cell types most associated with gene expression differences in the bulk data [58]). This revealed a common pattern in both mouse/treatment models (**Figure 7D**). With aging and tauopathy, increased gene expression signatures (i.e., of inflammation) were largely associated with glia, whereas decreased gene expression signatures were primarily associated with neurons—and in all cases these cell type-specific contributions to gene expression were at least somewhat reversed with Nanoligomer treatment. To confirm our RNA-seq data, we performed immunoblotting for several proteins whose gene expression levels changed (**Figure S4**), and we found similar patterns of protein expression for these targets. Also, given our findings above suggesting changes in glial cells and neuroprotection, we immunoblotted for CD68 (a marker of activated/reactive microglia [59]) and Brain-Derived Neurotrophic Factor (BDNF), a protein responsible for maintaining neuronal survival and growth [60] (**Figure 7E**). We observed reduced CD68 in Nanoligomer-treated animals, although this effect was not significant in old mice, but no treatment-associated changes in BDNF in either model. Although not definitive, these results suggest that Nanoligomer targeting of NF- κB/NLRP3 favorably influences several important biological processes associated with brain aging and tauopathy, particularly those involved in neuroinflammation/immune activation and neuronal health, and that these effects may be mostly driven by changes in glial cells, upstream of neuroprotection (i.e., because the inflammation-related gene expression patterns that changed with treatment mapped to these cell types in deconvolution analyses and neuron-intrinsic protective proteins like BDNF were unchanged).

### 2.7. Favorable peripheral changes in response to Nanoligomer treatment

Finally, current therapies meant to target neuroinflammation in the brain often have off-target effects [12, 61], and drug-induced hepatotoxicity is common following exposure to many bioactive compounds [62]. The liver is a key metabolic organ that processes such compounds and therefore a relevant target for therapies meant to help the brain [63]. To understand how Nanoligomer treatment affects the liver, we performed multiplex ELISAs on liver tissue from young, old, and old treated mice (**Figure 8A**). We found that several pro-inflammatory and immune-activating cytokines that increased with aging were reduced with Nanoligomer treatment, including IFN-Ɣ inducible protein (IP-10) and chemokine ligand 1 (CXCL1). Additionally, the anti-inflammatory cytokine interleukin 10 (IL-10) was reduced with aging, but this effect was reversed with Nanoligomer treatment. We also found that aging was associated with evidence of hepatocyte polyploidy and polymorphonuclear leukocytes in the liver, a key source of tissue damage due to inflammation [64], and these effects were attenuated with Nanoligomer treatment (**Figure 8B,C**). Furthermore, in Nanoligomer-treated old mice, these favorable changes in the liver were associated with a preservation of circulating (plasma) levels of fibroblast growth factor 21 (FGF-21), a hormone synthesized by the liver and linked with longevity and neuroprotection [65, 66]. Collectively, these results suggest that, in addition to their protective effects in the brain, Nanoligomers targeting NF-κB/NLRP3 are not hepatotoxic and may even favorably modulate several markers of inflammation and liver/peripheral health in old mice.

**Figure 8.**
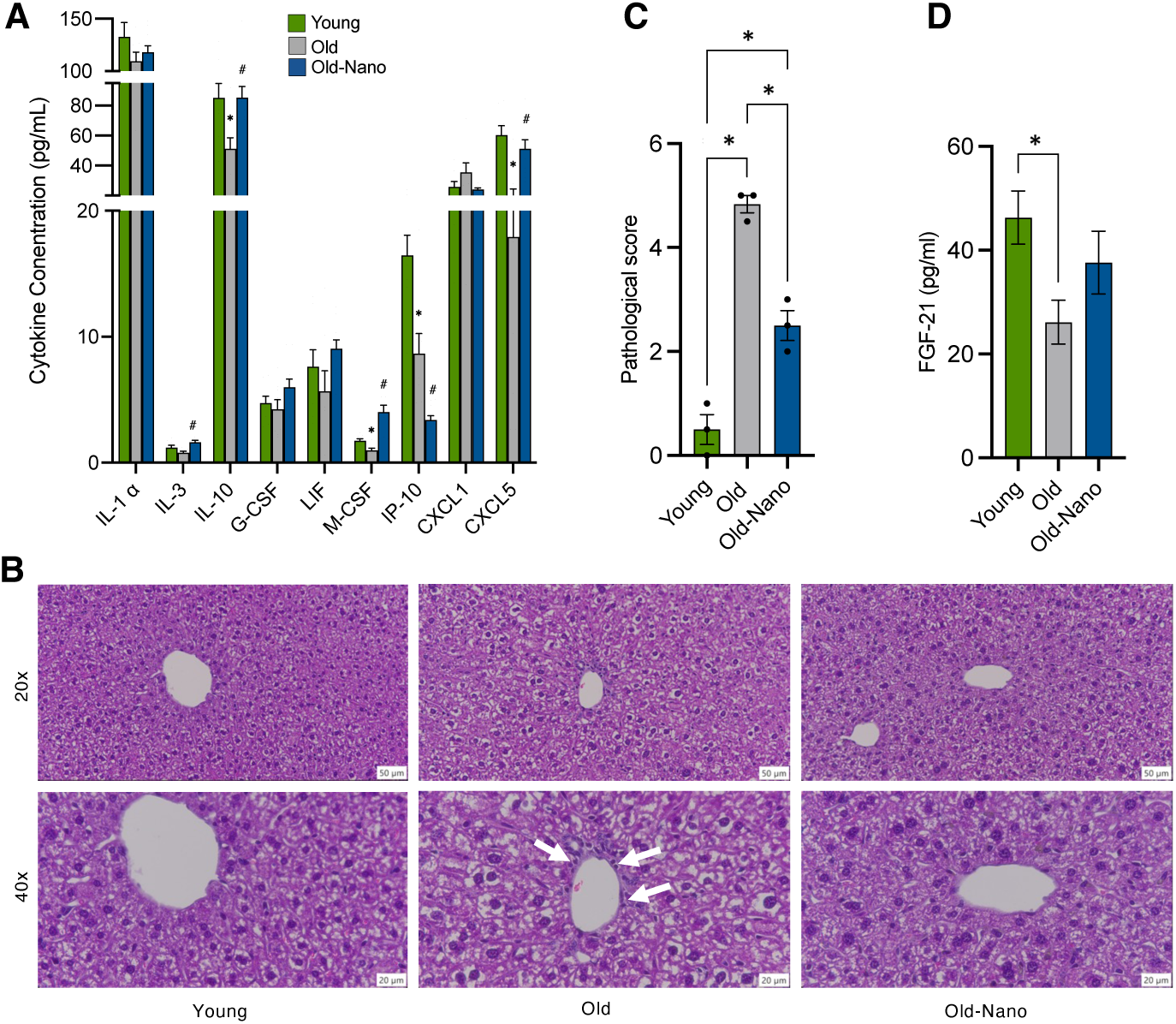
Favorable peripheral/systemic changes with long-term Nanoligomer treatment. **(A)** Cytokine concentrations in liver of young, old and old Nanoligomer-treated wildtype mice. N = 4/group; *p < 0.05 vs. young; ^#^p < 0.05 vs. old **(B)** Representative immuno-histochemistry images showing infiltrating immune cells at portal vein border (white arrows) and **(C)** quantification of immune cell infiltration in the same mice. *p < 0.01. **(D)** Plasma levels of fibroblast growth factor 21 (FGF-21) measured by ELISA in the same mice. Data analyzed via one-way ANOVA with group differences determined via Tukey’s post-hoc multiple comparison test. N = 6-11/group; *p < 0.01.

## DISCUSSION

Neuroinflammation impairs cognitive function and is a major mechanism of brain aging and AD [5], both of which are linked with tau pathology. As such, there is an urgent need to identify safe and effective treatments that target neuroinflammation [67]. Age- and tau-related neuroinflammation is characterized by immune activation, glial cell (microglia and astrocytes) reactivity, and the release of neurotoxic, pro-inflammatory cytokines [68, 69], and many of these events involve signaling via NF-κB and NLRP3. These two proteins are upstream mediators of immune activation and key components of a “central axis” of neuroinflammation [7, 22, 70]. For example, NF-κB regulates many inflammatory processes, including the activation of pro-inflammatory cytokines and the transcription of neuroinflammatory genes [22], and NLRP3 is a major component of the inflammasome that directly regulates multiple immune response pathways (e.g., complement/cytokine signaling) [71]. Our key finding is that simultaneous, targeted downregulation of both NF-κB and NLRP3 via Nanoligomers reduces neuroinflammation and improves cognitive function in mouse models of both aging and tauopathy. This may be an important, clinically relevant finding, as these small molecule therapeutics are well positioned for further pre-clinical and eventual clinical translation.

Several recent investigations have shown that separately targeting either NF-κB or NLRP3 reduces neuroinflammation and improves cognitive function in multiple mouse models [9, 36, 72, 73]. For example, an NF-κB peptide inhibitor was shown to reduce inflammatory cytokine expression and microglial activation in 5xFAD mice [11], and inhibition of NLRP3 with the small molecule MCC950 is reported to reduce inflammation and improve cognitive function in APP/PS1 mice [9]. However, although NF-κB and NLRP3 signaling pathways are interconnected [21], no studies have investigated the effects of simultaneously targeting both of these proteins in old and/or tauopathy mice. To investigate this idea here, we performed an initial proof-of-concept study and found that short-term treatment with Nanoligomers targeting NF-κB and NLRP3 caused reductions in target proteins in the hippocampus of old mice, consistent with our previous reports on this same cocktail both *in vitro* [15] and *in vivo* [18]. Then, to provide translational relevance to this finding, we performed a longer intervention (proportional to that which might be conducted in older adults), in which we treated both old and tauopathy model mice with NF-κB/NLRP3-targeting Nanoligomers for one month. Notably, we found that this intervention was safe, well-tolerated, and elicited no adverse neuromuscular effects (e.g., on grip strength or frailty) or in the liver, a key metabolic organ (i.e., despite the potential protective role of NF-κB in various settings). We also found that this treatment improved short-term recognition memory and reduced anxiety, which is often co-morbid with dementia [74]. These results are important, because cognitive decline is a major risk factor for mild cognitive impairment and AD [75], and our findings are consistent with data from other studies on NF-κB and NLRP3 in mouse models of aging and AD [9, 11]. We note that intraperitoneal Nanoligomer delivery (the mode of treatment in our study) is a translational limitation, but our group is currently developing alternative approaches for administering these compounds, including via intranasal inhalation and oral routes. Demonstrating the efficacy of such approaches will be an important next step in this line of investigation.

Importantly, numerous pro-inflammatory/cytokine signaling pathways contribute to neuroinflammation that drives declines in cognitive function [5, 69, 76]. To address this in the present study, we probed the hippocampus for multiple cytokines that have been linked with cognitive dysfunction, brain aging and tauopathy. We found that NF-κB/NLRP3-targeting Nanoligomer treatment reduced almost all of these cytokines, with PCA showing a marked reversal of overall cytokine patterns in both aged and tauopathy mice. Moreover, these cytokine profiles were highly (inversely) correlated with cognitive-behavioral function in all animals studied. Collectively, these results suggest that the mechanism by which Nanoligomer treatment improves cognitive function involves broadly reducing neuroinflammation. This idea is in agreement with our previous studies showing that Nanoligomer treatment reduces neuroinflammation and improves cognitive function in prion-disease mice (a model of severe neurodegeneration) [18], and with numerous other studies showing that targeting neuroinflammation improves cognitive function in mouse models of accelerated brain aging [77]. In future work, it will be important to identify the specific NF-κB/NLRP3-associated cytokine pathways that impact cognitive function most directly, as some cytokines (e.g., IL-10) can have both protective [78] and deleterious effects in the brain [79].

Beyond specific cytokine pathways, neuroinflammation is linked with protein pathology (including tau accumulation) and glial cell dysregulation in aging and AD [4, 80–82]. However, the order of these events is incompletely understood, and neuro-inflammatory signaling mediators like NF-κB and NLRP3 have been reported to both contribute to and be activated by pathology [36]. In our study, we observed several effects of NF-κB/NLRP3-targeting Nanoligomers that are consistent with this idea. First, we found that reduced neuroinflammation with NF-κB/NLRP3-targeting Nanoligomers was associated with reductions in total and phosphorylated tau species representative of both “early” and “later” aging/AD pathology in rTg4510 mice [39, 40]. Because aggressive, transgene-driven expression of tau occurs in rTg4510 mice at young ages, our findings may suggest that the treatment protected against tau-induced NF-κB/NLRP3 activation that further promotes pathology in these animals. This idea would be in line with other studies showing that tau seeding can be reduced by small molecule inhibitors of NLRP3 [35], and that pharmacologically targeting neuroinflammation, in general, reduces the burden of phosphorylated tau [83, 84]. Second, in aging wildtype mice, we found that NF-κB/NLRP3 Nanoligomer-induced inhibition of neuroinflammation was associated with evidence of reduced glial cell activation. In contrast to rTg4510 and other transgenic mouse models, aging wildtype mice do not develop tau or Aβ pathology. They do, however, develop glial cell-associated neuroinflammation, which has been linked with both age-related cognitive dysfunction and tau propagation [2, 5, 43, 69, 85]. Thus, our findings could suggest that NF-κB and NLRP3 also contribute to neuroinflammation regardless of pathology. Indeed, similar to previous studies showing that astrocytes and microglia become reactive/pro-inflammatory with aging [44, 49], here we found morphological and protein expression evidence of increased reactivity in both astrocytes and microglia of old mice—but NF-κB/NLRP3-targeting Nanoligomer treatment largely attenuated these effects. Together with our findings in tauopathy mice, these observations may suggest that NF-κB/NLRP3-associated neuroinflammation is a “proximal” target for enhancing cognitive function with aging and attenuating neurodegeneration (i.e., as depicted in Figure 1). Future, more comprehensive studies using NF-κB/NLRP3-targeting Nanoligomers are needed to confirm the order of these events, and/or to evaluate the treatment’s effects on other neuroinflammation-associated pathologies (e.g., Aβ aggregation). Such work may be particularly important given the notable limitations of the rTg4510 mouse model used here. For example, recent data suggest that pathology in these animals may be driven in part by off-target effects of the tau transgene (e.g., disruption of other genes important to brain function) [86]. In our data, these genes were modestly differentially expressed (average Log2Fold differences ∼-0.8) in transgenic animals, but their expression was not significantly altered with treatment and/or correlated with cognitive function metrics. This suggests that our overall findings may not have been driven by artifacts of the transgenic model but that the interpretation of our data should, perhaps, be taken with caution, and it underscores the need for similar studies in additional transgenic models (e.g., 5xFAD, APP/PS1 or humanized hTau mice).

Finally, given the protective effects of NF-κB/NLRP3-targeting Nanoligomers in both old wildtype and tauopathy mice, along with evidence that aging and tauopathy may intersect [87], we used transcriptomics (RNA-seq) to provide insight into conserved mechanisms of action. We found common transcriptome signatures of aging and tauopathy reflecting differences in neuronal health, neuroinflammation and immune activation, similar to ours and others’ observations in pre-clinical models of brain aging [88, 89] and aging/AD in humans [90]. However, these age- and tauopathy-associated transcriptome effects were largely attenuated with Nanoligomer treatment, and even reversed in some cases. Deconvolution analyses of these data also showed that increased gene expression patterns with aging and tauopathy were mostly associated with astrocytes and microglia, whereas decreased gene expression patterns were mostly associated with neurons. NF-κB/NLRP3-targeting Nanoligomers at least partially reversed these patterns in both models, which is consistent with the idea that the treatment largely affected glial cells (i.e., because the direction of gene expression changes in these cells tracked with that of inflammatory gene expression). Given the important role of NF-κB/NLRP3 in other cell types, additional multi-omics and/or perhaps single-cell analyses would be helpful in confirming these observations and more thoroughly evaluating the up- and down-stream mechanisms by which targeting NF-κB/NLRP3 protects cognitive function.

## CONCLUSIONS AND FUTURE DIRECTIONS

Our findings demonstrate that Nanoligomers specifically formulated to target NF-κB and NLRP3 are safe and well-tolerated, engage their target proteins in the brain, reduce neuroinflammation and improve cognitive function with both aging and tauopathy. Our molecular/transcriptomic data suggest overlapping mechanisms of action in these contexts, supporting the idea that NF-κB/NLRP3-associated neuroinflammation is a key target in aging and neurodegeneration. Future studies are needed to determine if this approach will work in larger animal models and/or humans, and to disentangle differential effects in short-term vs long-term treatment periods with varying dosages).

## METHODS

### Nanoligomer design and synthesis

Nanoligomers were synthesized by Sachi Bioworks as previously reported [13–18]. Briefly, Nanoligomers comprise an antisense peptide nucleic acid (PNA) conjugated to a gold nanoparticle for improved delivery and membrane transport. The PNA sequences, provided in [14, 15], were previously screened for solubility, self-complementarity, and off-targeting in the human genome. The PNA portions of the Nanoligomers (NLRP3 Sequence: CTTCTACTGCTCACAGG, NFKB Sequence: AGTGGTACCGTCTGCTA) were synthesized on a Vantage peptide synthesizer (AAPPTec, LLC) with solid-phase Fmoc chemistry. Fmoc-PNA monomers were obtained from PolyOrg Inc., with A, C, and G monomers protected with Bhoc groups. Following synthesis, the peptides were conjugated with gold nanoparticles and purified via size-exclusion filtration. Conjugation and concentration of the purified solution was monitored through measurement of absorbance at 260 nm (for detection of PNA) and 400 nm (for quantification of nanoparticles). The sequence of the synthesized PNA was confirmed using LC-MS [13].

### Animal husbandry

Older and younger male and female C57Bl/6J mice (aged 19 months and 8 months, respectively) were purchased from the National Institute on Aging aged rodent colony. Male and female rTg4510 mice and appropriate littermate controls (2 months of age) were purchased from the Jackson Laboratory (strain #024854). For the first part of the study (proof of concept to show that the Nanoligomer cocktail reduced both NF-κB and NLRP3 in the brain), cohorts were treated with Nanoligomer acutely (1x intraperitoneal injection; 150 mg/kg body weight) and then studied 16 hours later. For long-term treatment, mice were treated for one month (150 mg/kg body weight; 3x week). Mice were housed throughout all experiments at ∼18-23°C on a reverse 12 light/12 dark cycle. Fresh water and *ad-libitum* food (Tekland 2918; 18% protein) was routinely provided to all cages. Animals were consistently health checked by the veterinary staff at Colorado State University (CSU).

### Animal sacrifice and tissue collection

Mice were culled in a fed state in the late morning. After deep anesthetizing with isoflurane, ∼1 ml of blood was removed via cardiac puncture followed by cervical dislocation. The left hippocampus and a piece of left cortex tissue were removed, flash-frozen on dry ice, and stored at -80°C until further processing. The right hippocampus and right cortex were processed (paraffin-embedded) for immunostaining (details below).

### Immunoblotting

Frozen hippocampus samples were lysed in RIPA buffer containing 100 mM KCl, 40 mM Tris HCl, 10 mM Tris base, 5 mM MgCl_2_, 1 mM EDTA, and 1 mM ATP (pH 7.5), phoSTOP phosphatase inhibitors and cOmplete protease inhibitors (ThermoFisher). Sample protein concentration was determined using a BCA kit (ThermoFisher/Pierce), and 20 μg of protein per sample was separated by electrophoresis (Bio-Rad Criterion system), transferred to nitrocellulose membranes (Trans-Blot Turbo, Bio-Rad), and blocked in TBS-T with 5% Bovine Serum Albumin (BSA). Primary antibodies were purchased from ABclonal and included phosphorylated nuclear factor kappa B (p-NF-κB, 1:1000 dilution); nuclear factor kappa B (NF-κB, 1:1000); NLR Family Pyrin Domain Containing 3 (NLRP3, 1:1000); total tau (1:1000); p-tau S396 (1:1000); p-tau T205 (1:1000); Cluster of Differentiation 68 (CD68, 1:1000); Indoleamine 2,3-dioxygenase 1 (IDO1, 1:1000); Interleukin 17 Receptor E (IL17RE, 1:1000); and Brain-Derived Neurotrophic Factor (BDNF, 1:1000). GAPDH (1:2000) was purchased from Novus Biologicals. Membranes were incubated in primary antibody (TBS-T in 5% BSA) for 24 hours at 4°C, followed by HRP-conjugated secondary antibodies (TST-T in 5% milk; 1:5000; Cell Signaling) for 1 h. Membranes were then incubated in ECL, imaged on a ProteinSimple Fluor Chem imaging system, and quantified with ImageJ.

### Cognitive-behavioral and functional testing

Novel Object Recognition: For NOR testing, mice were assessed for short-term memory performance using no habituation phase [91]. Briefly, mice were placed in an opaque arena (50 cm x 50 cm) with two identical objects (tower of Legos) placed 8 cm from each side wall. After an inter-trial interval of two hours, mice were returned to the arena, but one of the old objects was replaced with a new object of similar shape and color (cell culture flask filled with multi-colored sand). Trials ended once 20 s total exploration time was reached, or 10 min had elapsed. If mice were unable to meet the exploration criteria of 20 s or 10 min, they were excluded from the analysis. Recognition index was calculated as novel object exploration time divided by 20 s [91].

Elevated Plus Maze: Elevated Plus Maze testing was performed as previously described [90, 92] and consisted of a cross with four arms (67 cm in length; 2 exposed arms, 2 enclosed arms). Mice were placed in the center of the maze facing an open arm and allowed to freely explore for 5 min. The amount of time in open arms (a measure of anxiety-like behavior) was recorded and analyzed with AnyMaze software.

Grip Strength: The grip strength test was performed as previously described [93]. Mice were tested using the DST-110 Digital Force Gauge (Maze Engineers). A total of 5 trials were performed per mouse and the highest and lowest values were discarded for analysis. The remaining 3 force values were averaged for each mouse. Data are presented as g force/g body weight.

Frailty Testing: Mouse frailty scores were assessed using an established scale [94]. Briefly, trained investigators blindly scored mice on a 31-point scale, which included measurements of health in the following areas: integumentary, physical/musculoskeletal, vestibular/auditory, ocular/nasal, digestive/urogenital, and respiratory systems. Higher scores indicate worse outcomes, while lower scores indicate better outcomes.

### Multiplexed ELISAs

Brain homogenates from all mice were assessed with a 36-plex Procartaplex cytokine/chemokine panel (ThermoFisher) as previously reported [15, 16], which included the following cytokines and chemokines: ENA- 78 (CXCL5), Eotaxin (CCL11), GRO-α (CXCL1), IP-10 (CXCL10), MCP-1 (CCL2), MIP-1α (CCL3), MIP-1β (CCL4), MIP-2α (CXCL2), RANTES (CCL5), G-CSF (CSF-3), GM-CSF, IFN-α, IFN-γ, IL-1α, IL-1β, IL-2, IL-3, IL-4, IL-5, IL-6, IL-9, IL-10, IL-12p70, IL-13, IL-15/IL-15R, IL-17A (CTLA-8), IL-18, IL-22, IL-23, IL-27, IL-28, IL-31, LIF, MCP-3 (CCL7), M-CSF, and TNF-α. For systemic effects (Figure 7), separate individual ELISAs were used. Briefly, 25uL of 10 mg/ml hippocampus, cortex, liver homogenate, or serum was processed using standard ELISA techniques and analyzed on a Luminex MAGPIX xMAP instrument. Standards for each cytokine/chemokine were used with 1:4 dilutions (8-fold dilutions) and background, controls and sample concentrations were determined from a standard curve using Five Parameter Logistic curve fit/quantification.

### Immunohistochemistry and immunofluorescence

Brain and liver tissues were fixed in 10% neutral buffered formalin at room temperature for at least 48 hours. Brain samples were processed on a Leica TP1020 Automatic Benchtop Tissue Processor and embedded in paraffin wax (Cancer Diagnostics). For immunohistochemistry analyses of phosphorylated tau, paraffin-embedded tissue was sectioned at 4 µm, deparaffinized, rehydrated, sodium citate treated, and blocked in Tris A/2% donkey serum (Jackson ImmunoResearch), then incubated overnight in the following antibodies: phospho-tau (Thr181; 1:400 dilution; Invitrogen) and phospho-tau (Thr231; 1:400; ABClonal). Wash steps were performed using 2% bovine serum albumin and 2% Triton-X in 1 M TBS, and an ABC HRP peroxidase detection kit (Vector Laboratories) and ImmPACT DAB Substrate Peroxidase (HRP) Kit (Vector Laboratories) were used for a chromogen. Slides were counterstained with hematoxylin (ThermoFisher), secured with a coverslip in mounting medium and stored at room temperature until imaging. Whole tissue images were taken on an Olympus BX53 microscope with an Olympus DP70 camera using an Olympus UPlanSApo 20x objective (N.A. = 0.75). Representative images were taken using an Olympus BX53 microscope with an Olympus DP70 camera using an Olympus UPlanFL N 40x objective (N.A. = 0.75). To quantify and select the phosphorylation of tau-positive stained cells the manual threshold on the Count and Measure function of Olympus CellSens software (v1.18) was used. For immunofluorescence analyses of GFAP, IBA1 and glial cell process length and branching, sections were incubated overnight in primary antibodies: IBA1 (1:50 dilution; Abcam) and GFAP (1:100; Dako). Slides were washed with TBS and incubated for 1 h in the dark with AlexaFluor 555 or 647 (Invitrogen) secondary antibodies at 1:500 dilution and 2% normal donkey serum, then washed and incubated in Hoechst dye (1:2000; ThermoFisher) for 3 min. Slides were mounted with Prolong Gold Antifade mounting media (ThermoFisher) and stored in the dark at 4°C prior to imaging. Images were captured using an Olympus BX63 fluorescence microscope equipped with a motorized stage and Hamamatsu ORCA-flash 4.0 LT CCD camera using a 40x Olympus X-Apochromat air objective (N.A. = 0.80). For astrocyte/microglial process length and number analyses, four regions between the dentate gyrus and CA1-CA3 region of the hippocampus were imaged at 40x. To identify/quantify processes and branches, skeletonization of GFAP+ astrocytes and IBA1+ microglia was performed using IMARIS software as previously described [95] to trace and quantify stained filaments. For histology analyses of liver, tissues were sectioned on a ThermoFisher HM 325-2 microtome at 5μm thickness and mounted on positively charged glass slides (Superfrost Plus, Cancer Diagnostics). Liver sections were deparaffinized and stained with hematoxylin and eosin (Cancer Diagnostics), and whole tissue images were taken for analysis using an Olympus DP70 camera and an Olympus UPlanSApo 20x objective. Livers were scored for pathological infiltration of immune cells specifically around the portal triad, and for overall hepatocyte morphology by a trained animal pathologist.

### RNA sequencing and transcriptomic analyses

Transcriptome (RNA-seq) and gene expression analyses were performed on whole hippocampus via methods previously described [89]. Frozen samples (n=6/group; 3M and 3F) were transferred into Trizol (ThermoFisher) and homogenized on ice (SP Bel-Art ProCulture Micro-Tube Homogenizer System; VWR Scientific). RNA was isolated using an RNA-specific spin column kit (Direct-Zol, Zymo Research) and treated with DNase to remove genomic DNA. Poly(A)-selected libraries were generated using magnetic oligo dT beads (ThermoFisher) and Illumina TruSeq kits. Libraries were sequenced on an Illumina NovaSeq 6000 platform to yield >40 M 151-bp paired-end FASTQ reads/sample. Reads were trimmed and filtered with the fastp program [96], then aligned to the mm10 *Mus musculus* genome using the STAR aligner [97]. Differential gene expression was analyzed with DESeq2 [98] and genes/transcripts with BaseMean read counts of <10 were filtered and removed for analyses [99, 100]. Differentially expressed genes were analyzed for gene ontology (GO) enrichment using the Metascape program [101], and heatmaps were constructed with GraphPad Prism software. Deconvolution analyses of bulk RNA-seq data were performed as previously described [88] using a published protocol and R script [58]. Briefly, single-cell RNA-seq data on mouse hippocampus (SMART-seq, 2019; 10x-SMART-seq taxonomy, 2020) were downloaded from the Allen Brain Map portal [102], and cells from the hippocampus, including astrocytes, endothelial cells, microglia, oligodendrocytes and neurons, were sub-setted for analysis. Normalized DESeq2 gene counts for old vs. young mice, old Nanoligomer-treated vs. old, rTg4510 vs. LM Controls, and rTg4510 Nanoligomer-treated vs. rTg4510 comparisons in bulk RNA-seq data were filtered for the top 250 genes by p-value, and then sorted by Log2Fold change. Sorted gene lists were scaled to the single-cell RNA-seq dataset, normalized expression of the same genes in the single-cell dataset was obtained and used for dimensional reduction via principal component analysis and hierarchical clustering/heat mapping to obtain cell-type specific contributions to differences in the bulk RNA-seq comparisons, and cell types associated with each cluster were confirmed using CellMarker 2.0 [103].

### Statistics

One-way ANOVA with Tukey’s *post-hoc* testing was used to assess differences among groups in protein/cytokine levels and measures of cognitive function, as well as glial cell morphology, and simple linear regression (Pearson correlations) were used to relate cytokine levels to cognitive function data. A non-parametric ANOVA equivalent (Kruskal-Wallis test) was used for small sample sizes with non-normally distributed data. All data were analyzed and presented using GraphPad Prism software. Differentially expressed genes were detected using DESeq2, and genes were sorted by p-value and Log2Fold change to construct volcano plots and heatmaps, respectively. The Metascape program was used for GO and TRRUST analyses [101].

## Author contributions

D.W. designed the study, wrote the paper, generated/analyzed data, conducted behavioral experiments, and generated/analyzed RNA-seq and immunoblot data; S.J.R. generated and analyzed immunofluorescence/immunohistochemistry data; S.C.O, T.E., V.S.G., and A.C. generated and analyzed cytokine/protein data; P.N. designed the study, provided conceptual insight, and generated/analyzed data; J.A.M. designed the study and generated/analyzed immunofluorescence/immunohistochemistry data; T.J.L. designed the study, wrote the paper, generated/analyzed data, provided conceptual insight, and provided funding. All authors read and approved the final manuscript.

## Funding

This project was supported in part by NIH awards AG078859, AG070562 and AG060302 (T.J.L.) and AG069361 (D.W.), NASA SBIR Awards 80NSSC22CA116, 80NSSC23CA171 and National Academy of Science Health Longevity Catalyst Award (P.N.), and by intramural (startup) funding from Colorado State University (T.J.L.).

## Data Availability

The RNA-seq data generated in this study are available on the Gene Expression Omnibus website under accession number GSE251888. All additional data are included in the data supplement, and raw data will be made available upon request to the corresponding author (T.J.L).

## Declarations

### Ethics approval and consent to participate

This protocol was approved by IACUC #1441 and Laboratory Animal Resources (RRID: SCR_022157) staff at Colorado State University.

### Consent for publication

Not applicable.

### Competing interests

S.S, V.G, A.C., and P.N. work at Sachi Bio, a for-profit company, where the Nanoligomer technology was developed. A.C. and P.N. are the founders of Sachi Bio, and P.N. has filed a patent on this technology. All other authors declare no competing interest.

## Supplementary Figures

**Figure S1.**
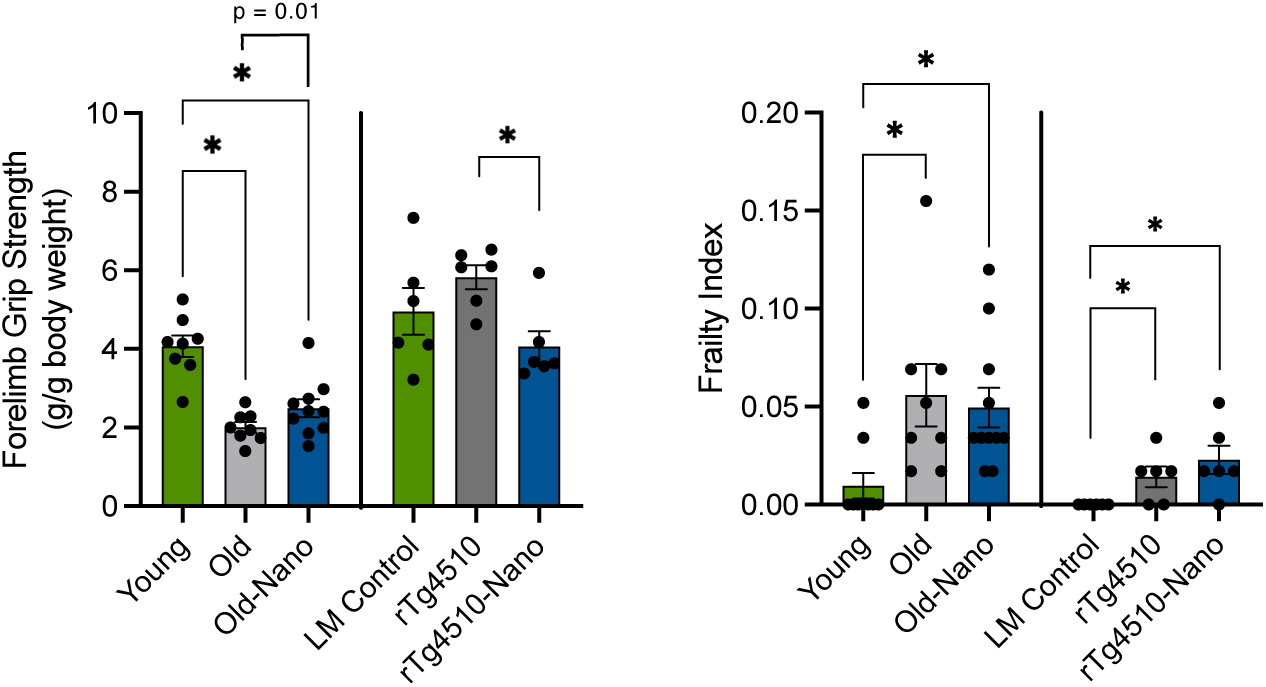
Nanoligomers targeting NF-κB and NLRP3 increase grip strength in old wildtype mice and have no adverse effects on overall physical health/frailty. Left: Forelimb grip strength in young, old and old Nanoligomer-treated wildtype mice, as well as littermate (LM) controls, rTg4510 tauopathy and rTg4510 Nanoligomer-treated mice. Right: Frailty Index in the same animals. N = 6-11/group; *p < 0.05.

**Figure S2.**
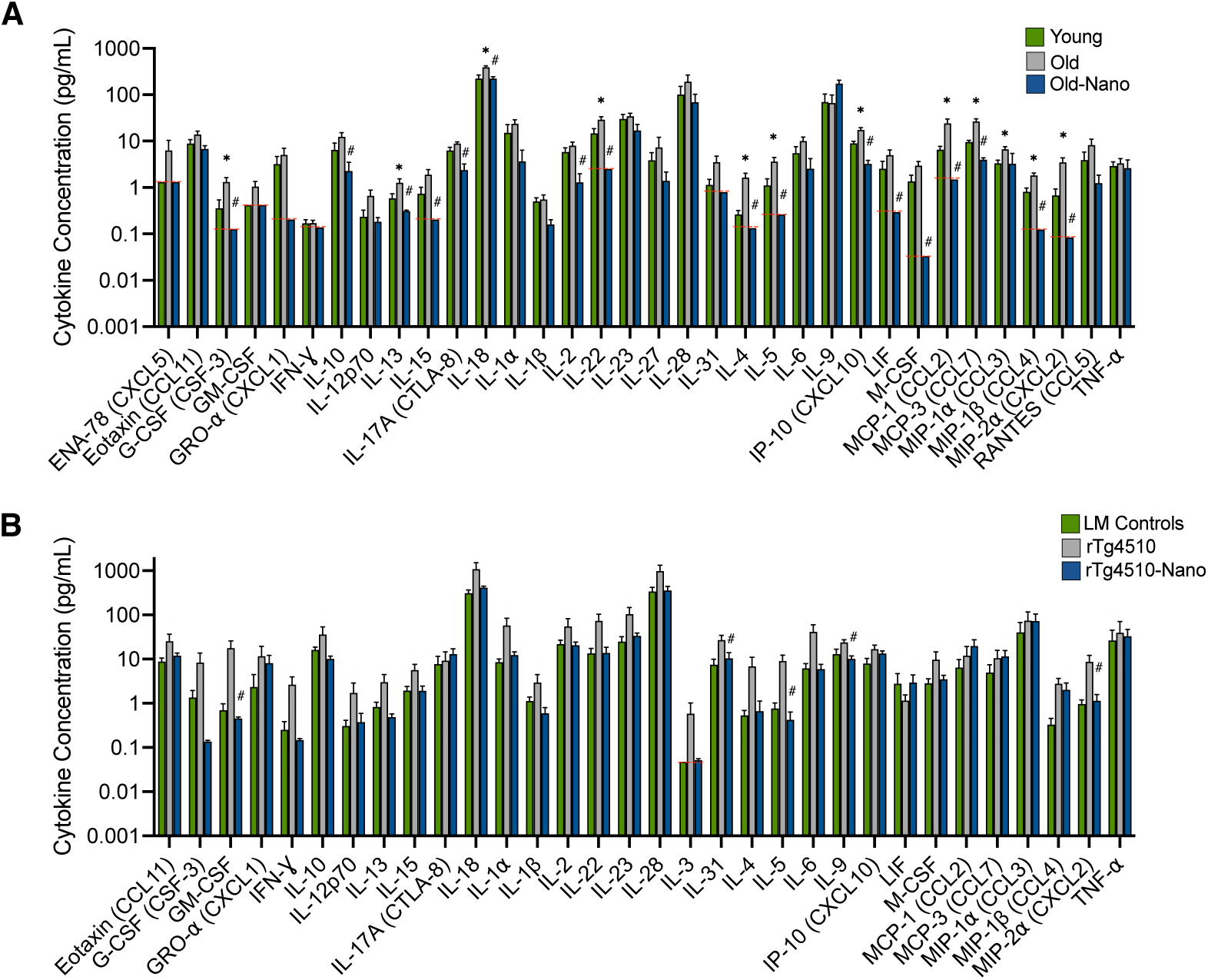
Nanoligomers targeting NF-κB and NLRP3 reverse age- and tauopathy-related increases in many cytokines in the brain. Multiplex ELISA analyses of prefrontal cortex tissue from young, old and old Nanoligomer-treated wildtype mice, as well as littermate (LM) controls, rTg4510 tauopathy and rTg4510 Nanoligomer-treated mice. N = 6-11/group; *p < 0.05 vs. young/LM; ^#^p < 0.05 vs. old/rTg4510; red lines represent limit of detection.

**Figure S3.**
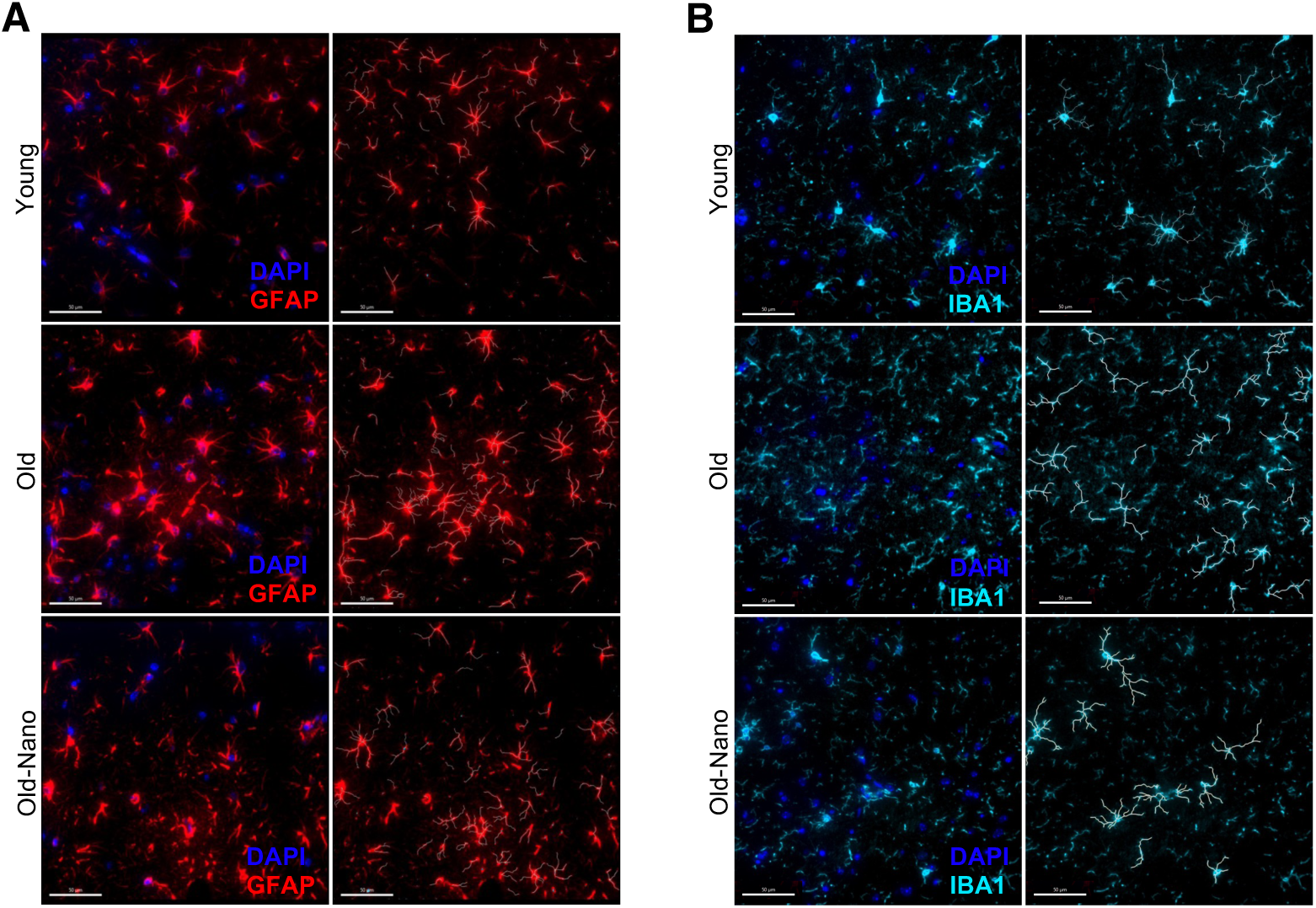
Nanoligomers targeting NF-κB and NLRP3 modulate glial cell morphology in old mice. Representative images from Figure 5 (left in each panel) and corresponding images (right in each panel) showing skeletonization (white lines) for assessing morphology in **(A)** astrocytes stained for GFAP and **(B)** microglia stained for IBA1.

**Figure S4.**
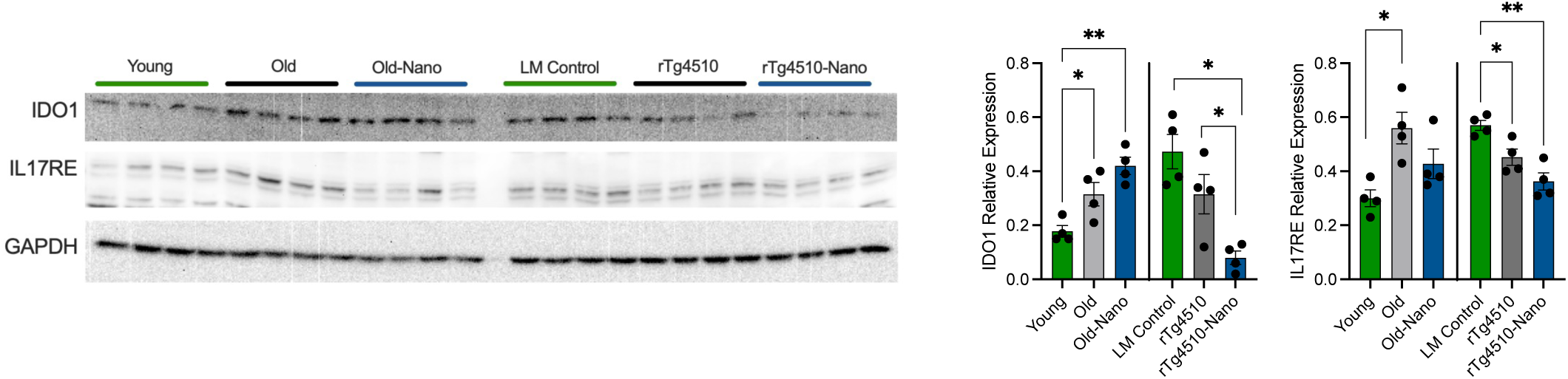
Immunoblotting confirmation of hits in RNA-seq data. Raw immunoblots and quantifications showing protein levels that track with RNA-seq gene expression levels for Indoleamine 2,3-dioxygenase 1 (IDO1) and Interleukin 17 Receptor E (IL17RE), both of which have also been linked with differences in cognitive function, in young, old and old Nanoligomer-treated wildtype mice, as well as littermate (LM) controls, rTg4510 tauopathy and rTg4510 Nanoligomer-treated mice. *p < 0.05; **p < 0.01. Note: same blot/GAPDH loading controls as in Figure 7.

## Notes

### Competing Interest Statement

Co-authors S.S, V.G, A.C., and P.N. work at Sachi Bio, a for-profit company, where the Nanoligomer technology was developed. A.C. and P.N. are the founders of Sachi Bio, and P.N. has filed a patent on this technology. All other authors declare no competing interest.

### Summary of Updates

Additional data and analyses have been added, along with text describing and discussing them.

